# Antibacterial 3D-printed PMMA/ceramic composites

**DOI:** 10.1101/2021.10.11.463892

**Authors:** Elia Marin, Mikiya Mukai, Francesco Boschetto, Thefye P. M. Sunthar, Tetsuya Adachi, Wenliang Zhu, Alfredo Rondinella, Alex Lanzutti, Narisato Kanamura, Toshiro Yamamoto, Lorenzo Fedrizzi, Giuseppe Pezzotti

## Abstract

Following the rising interested on 3D-printed technologies, this research explores the possibility to use stereo-lithography to 3D print PMMA resins reinforced with up to 15% in weight of antibacterial ceramic powders. Three different reinforcements were tested, following previous literature data: aluminum nitride, titanium oxide and barium titanate.

Between the three powders, the most uniform dispersion was achieved using aluminum nitride. Initial screenings with mixed and cured composite resins showed that between the three composite materials, only aluminum nitride or barium titanate PMMA showed a clear antibacterial effect when compared to the pristine reference, with aluminum nitride being the most effective against *E. coli*. When 3D printed using stereo-lithography, the composite containing aluminum nitride showed an even higher degree of dispersion and comparable antibacterial effects. Moreover, aluminum nitride reinforced PMMA resins showed good mechanical properties, comparable to the basic resin, and could be further strengthened by a standard post-curing process.

## 1. Introduction

Poly-methacrylate is one of the few groups of polymers that could find an application as structural materials in the orthopedic and orthodontic fields, the others beings polyethylene and poly-ether-ether-ketone. When compared to the other two, poly-methacrylate has usually a lower mechanical strength, which is however more than compensated by the higher flexibility of use.

Low production costs, chemical and photo/chemical durability and good light transmittance in the visible range made PMMA the most successful polymer in the field of disposable contact lenses, where it has been applied since 1950s [1].

In dentistry and orthopedics, thermosetting PMMA-based resins have been successfully used for decades [2]. The most common application in these fields are as “bone cement”, acting as a filler for bone losses, or as a “glue” between bone and prosthetic/orthopedic implants [3]. Orthodontic implants made of PMMA cements can also be used as long term bone-substitutes in load bearing applications, if the load is not excessive [4].

The use of PMMA in biomedical applications has been approved mainly because of its lack of toxicity, chemical stability over time and good mechanical properties. PMMA usually doesn’t cause inflammatory reactions with tissues and, even if not bio-tolerated it is considered sufficiently bio- inert.

Like many other polymers, PMMA resins can also be combined with other materials or molecules in order to further enhance their properties and produce composites. A few notable examples are nanotubes reinforced PMMA, which showed higher mechanical strength and electrical conductivity [5], ZrO_2_ reinforced PMMA, which could withstand higher friction contact loads, resulting in lower wear rates when compared to pristine references [6] and zincite (ZnO) functionalized PMMA, applied in the optical field for its high refractive index [7].

For biomedical applications, the bio-inertness of PMMA is often considered the most severe limit for its applicability. Bio-tolerated or even bio-active materials are nowadays preferred in most tissue regeneration applications, as bio-inert materials might result in a slower recovery or facilitate the penetration of bacteria in the open wound. Moreover, patients with ongoing bone-related conditions, such as osteoporosis, might not be able to recover from surgery without a bioactive support. PMMA has been functionalized with a wide array of drugs and particles to both enhance bioactivity and reduce risk of surgical site infections. ZrO_2_ particles have been successfully used for esthetical dental restoration [8], bioglass to stimulate bone tissue formation [9], antibiotics [10] or silver particles [11] to prevent infections.

Nowadays, acrylic resins are one of the most common polymeric material used in both commercial and hobbyist 3D-printing, in particular in stereolitography (SLA) and Digital Light Process (DLP). Even if much slower than most conventional manufacturing methods, stereolitography (SLA) is considered a promising additive manufacturing techniques both for polymeric [12] and for ceramic materials [13], in particular for custom-made dental orthodontic devices [14]. When compared with other techniques, such as extrusion printing [15], SLA results in better surface finishing, superior mechanical strength and higher geometrical precision [16].

Most stereo-lithographic additive processes require three additional steps after printing: a rinse to remove excesses of liquid resin, the removal of extra support material and curing at a specific wavelength (usually UV-A).

Due to the versatility of the process, SLA can be easily adapted to produce PMMA-based composite materials, and various attempts have been reported in literature. The most common reinforcements for PMMA are graphene particles [17] and metal and metal oxides [18], but so far these materials found little if any industrial application.

In this research, we explore the anti-bacterial effects of PMMA composites consisting of 5%, 10% or 15% of three different reinforcing ceramic particles, namely rutile, barium titanate and aluminum nitride. Barium titanate was selected as a potential candidate after showing promising antibacterial effects when used as a reinforcement in other polymeric matrices [19, 20], while aluminum nitride was already successfully applied in PMMA-based coatings [21]. After a preliminary characterization the best performing candidate (AlN) was selected and used to produce a complex 3D printed part using stereo-lithography. The parts, construction blocks similar to those used as toys for kids, were then tested *in vitro* against *Escherichia coli*, showing a good protection against bacteria adhesion.

The contents of this research are summarized in Figure 1.

**Figure 1:**
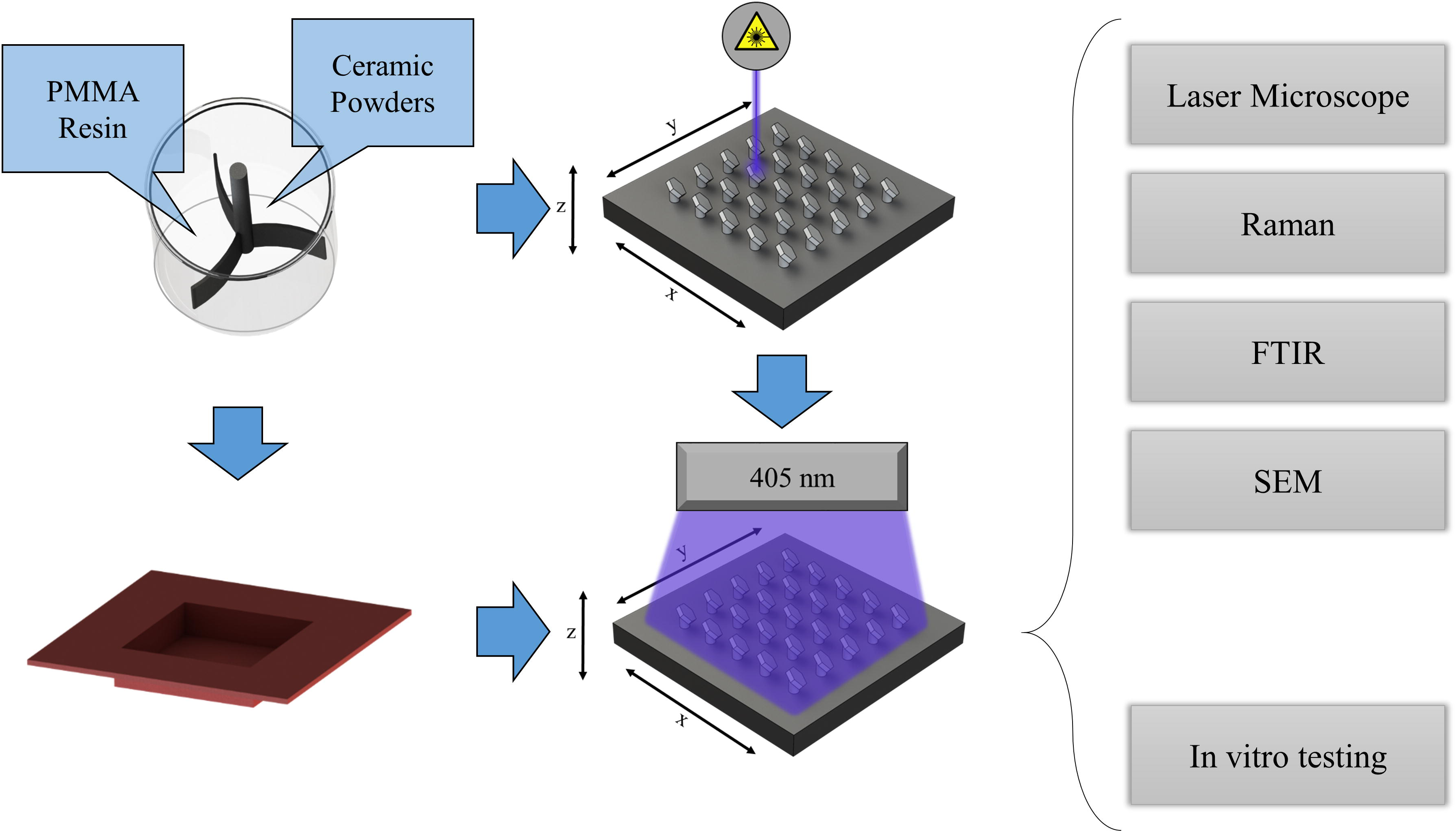
Schematic of the research: composite polymer preparation, curing and testing

## 2. Experimental

### 2.1 Ceramic powders

Three different ceramic powders, namely aluminum nitride (AlN), barium titanate (BaTiO_3_) and rutile (TiO_2_) were acquired from an industrial producer (Sigma Aldrich, Missouri, United States). The declared average powder diameter was about 5 μm for AlN, 1 μm for BaTiO_3_, 1 μm for TiO_2_. The diameter of the aluminum nitride powder was chosen in order to reduce surface energy and thus oxidation in humid air environments.

### 2.2 Preliminary sample preparation

The composite resins were obtained by mechanical mixing ceramic powders and PMMA resin using an impeller at a rotating speed of 120 rpm, for 60 minutes. The resin (Clear Photoreactive Resin, Formlabs, Somerville, Massachusetts, US) is composed of methacrylated oligomer (about 75%), methacrylated monomer (about 25%) and diphenyl (2,4,6-trimethylbenzoyl) phosphine oxide (≤ 1%) as a photo-initiator.

Resins containing 5, 10 and 15% in weight of ceramic reinforcement were then poured into silicon molds of 40 x 50 x 2 mm, heated up to 150 °C for 1 hour, cooled down and post-cured under a 405 nm UV light for 20 minutes. This treatment was used to assure the complete solidification of the resin samples used during the preliminary testing.

### 2.3 Stereolitographic printing

3D-printed samples were produced using a Form2, a commercial stereolithographic 3D printer with a 405 nm laser source (Formlabs, Somerville, Massachusetts, US). The laser positioning is controlled by two X-Y step motors with a maximum theoretical resolution of 150 µm, while the stage position (Z axis) has a nominal layer thickness resolution of 25 µm.

To reduce the need for additional supports and optimize the geometrical tolerances, samples were printed at a suggested tilting angle of 45° (Figure 1a). The stereolithographic process was performed at 31 °C, with a set theoretical layer resolution of 50 µm.

In this process, only aluminum nitride could be printed correctly, while both barium titanate and rutile produced incomplete and distorted parts and were therefore excluded from the manuscript.

For 3D printing, two different sample designs were used: a short, flat dog-bone shaped with a thickness of 1 mm, a gage width of 6 mm and a gage length of 20 mm, used for the mechanical testing, and kid construction blocks of about 31.8 x 15.8 x 9.6 mm with two sets of four holes and studs and used for final antibacterial characterizations and tolerance checking.

The samples were then post-cured at 60 °C for 20 minutes (Form Cure, Formlabs, Somerville, Massachusetts, US, Figure 1c) and washed in isopropyl alcohol (KT Chemicals Co., Ltd., Osaka, Japan) for 20 minutes (Form Wash, Formlabs, Somerville, Massachusetts, US).

### 2.3 Characterization

#### 2.3.1 Scanning Electron Microscopy

A SM-700 1F Scanning Electron Microscopy (SEM) (JEOL, Tokyo, Japan) was used to acquire images at magnifications ranging from 50× to 50,000×. SEM images were used to observe the morphology of the ceramic powders and estimate their average diameter. The samples were sputtered with a layer nanometric of platinum before being observed at an accelerating voltage of 15 kV. Observations were also performed on the composite resins, coupled with Electron Dispersive X-ray Diffraction (EDXS) analyses obtained using an external module (X-Max EDS system, Oxford Instruments, Oxford, UK).

#### 2.3.2 Laser Microscopy

A VKX200K series laser microscope (Keyence, Osaka, Japan) was used to obtain low magnification images of large areas of the samples. The instrument could obtain both optical and topographical information on the surface morphology, with magnifications ranging from 10× to 150× and a numerical aperture between 0.30 and 0.95. The stage positioning could be controlled using by two step motors, while the autofocus function controlled the distance between the lenses and the surface of the sample, in the z range, allowing to perform automated focus stacking.

#### 2.3.3 Raman Spectroscopy

Raman spectra were collected using T-64000 spectrometer (Horiba, Kyoto, Japan) based on a triple monochromator and equipped with a 1024 × 1024 channels charged coupled device (CCD) detector.

The acquired spectra were then elaborated using a commercially available software (LabSpec, Horiba, Kyoto, Japan and Origin, Originlab Corp., Northampton, MA, US).

The excitation source was a 514 nm Ar-ion laser (Stabilite, Spectra-Physics, Massachusetts, United States) operating at the nominal power of 80 mW. By using a 100x magnification objective lens, the probe area could be reduced to a diameter of about 1 µm. The later movement of the stage was controlled by two step motors with a resolution in the order of 1 µm. All spectra were acquired and elaborated using a dedicated software (Labspec 5.0, Horiba, Kyoto, Japan).

Raman imaging was obtained using a dedicated Raman microscope coupled with a spectrometer (RAMANtouch, Nanophoton Co., Ltd., Osaka, Japan). The excitation source was a green light with a wavelength of 532 nm, produced by a diode laser. The instrument operated at a maximum nominal power of 200 mW. The power was set to an appropriate value by preliminary adjustment of neutral density filter. The microprobe used lenses ranging from 5x to 100x magnifications with numerical apertures from to 0.5 to 0.23.

#### 2.3.4 Fourier Transformed Infra-Red Spectroscopy

Fourier Transformed Infra-Red Spectroscopy (FTIR) was performed using a dedicated spectrometer (FT/IR-4000 JASCO, Tokyo, Japan) operating at room temperature. The instrument was equipped with a Michelson 28 deg interferometer with corner-cube mirrors, and was able to cover a theoretical spectral range between 250,000 and 5 cm^-1^.

The fixed aperture size was 200×200 µm^2^ and the acquisition time was set to 30 s and 10 repetitions. The instrument operated using a dedicated software (Spectra Manager, JASCO, Tokyo, Japan). All signal post-processing, baseline removal and deconvolution were performed using Origin (Originlab Corp., Northampton, MA, US).

#### 2.3.5 Mechanical testing

The tensile testing, performed on the dumbbell shaped 3D printed samples, was achieved using a MCT 2150 Desktop Tensile-Compression Tester (AND Discover Precision, Tokyo, Japan) at a constant displacement rate of 10 mm/min. During testing, load and elongation were logged to a computer terminal, at a rate of 10 measurements/second. The tensile properties were determined as an average from 6 independent tests for each concentration, before and after UV post-curing.

### 2.4 In vitro testing

The culture medium was prepared in accordance with a previous protocol [22]: a phosphate-buffered saline solution (PBS) was selected as a medium due to the ion concentrations comparable to those of blood. The medium was then supplemented with 7.0% of glucose, which acted as an energy source, and plasma (human, 10%), used as a source of fundamental proteins.

Samples of the medium were inoculated with a strain of *Escherichia coli* (*E. coli*, ATCC® 25922™) and the solutions were then agitated on heated plate shaker (SBT1500-H, Southwest Science, Hamilton, NJ, USA) at 37°C and 175 rpm for one day. After incubation, the bacteria could reach a concentration of about 10^5^ cells/mL.

Then, each sample was inoculated with the bacterial solution within individual well plates. The well plates were subsequently placed on the shaking incubator for 12 h or 24 h at 37°C and 120 rpm.

#### 2.4.1 Colony counting

Two-hundred microliters of the bacterial and fungal solutions were dropped onto the surface of the samples (3 cm × 3 cm) and covered with parafilm. The samples were then incubated for 24 h at 37°C. After the incubation period, the samples were washed with 5 mL of sterilized phosphate-buffered saline (PBS) (Fujifilm WakoCo, Osaka, Japan). Then, one-hundred microliters were taken out from the mixture and placed into the microcentrifuge tube for 10-fold serial dilution from 10^−1^ to 10^−7^ by PBS. Finally, one-hundred microliters of each dilution were spread on an agar plate and incubated at 37° C for 24 h, which was followed by counting the number of colonies of the respective dilution. The CFU/mL was calculated using the formula: CFU/mL = (no. of colonies × dilution factor) × 10. Then, the CFU values were multiplied by the polyethylene surface and divided by the area of the parafilm used.

#### 2.4.2 WST

To observe and compare the bacterial metabolism after exposure of 12 and 24 hours, the samples were analyzed using colorimetric assay (Microbial Viability Assay Kit-WST, Dojindo, Kumamoto, Japan). This technique is based on the employment of a colorimetric indicator (WST-8) which, upon reduction in the presence of an electron mediator, produced a water-soluble formazan dye. The amount of the formazan dye generated was directly proportional to the number of living microorganism. Solutions were analyzed using microplate readers (EMax, Molecular Devices, Sunnyvale, CA, USA) upon collecting the OD value related to living cells.

#### 2.4.3 Fluorescence testing

3D printed parts were observed by UV light after in vitro testing for 24 hours with E. coli and subsequently being marked with a fluorescent probe (Gene 6 *E. coli* transformation kit, Nippongene, Toyama, Japan). Images were then collected using a dedicated, custom-made camera based on a 12 MP, f/1.7, 26mm (wide), 1/2.55”, 1.4µm, dual pixel PDAF, OIS.

### 2.5 Statistical significance

For statistical purposes, all spectra have been acquired in 10 different locations and all biological and mechanical tests were repeated on 5 samples for each concentration of reinforcing particles. Statistical significance has been addressed by two-way ANOVA analysis of variance. Statistically significant results (p < 0.05) have been marked with an “*”.

## 3. Results

Figure 2 shows scanning electron microscope images of the three different ceramic powders used in this research. It can be observed that each powder has a specific grain shape and granulometry, with rutile TiO_2_ (c) being the finest and AlN (b) being the coarser. Barium titanate (a) and rutile (c) also share a similar morphology, with polyhedral grains with an average size of about 1 μm. In the case of oxide-based powders, a smaller size was chosen in order to ideally achieve a better dispersion in the PMMA matrix. For aluminum nitride, a coarser powder size (5 μm) was selected in order to limit surface oxidation due to the high surface energy of the nitride particles.

**Figure 2:**
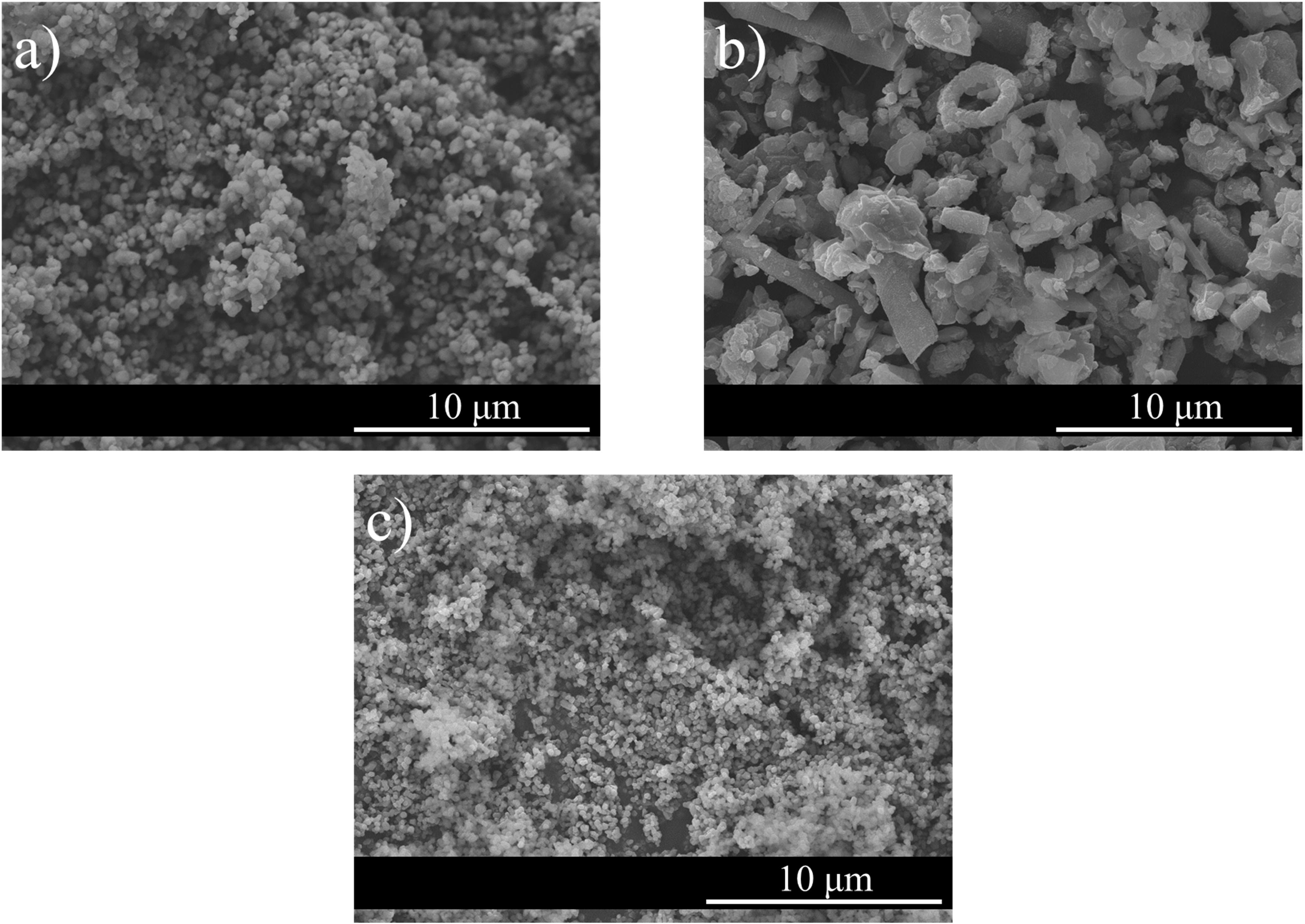
Scanning Electron Microscope ceramic powders: a) barium titanate, b) aluminum nitride and c) rutile

Figure 3 shows SEM images of the different composite materials containing 15% of reinforcing particles, compared to the pristine PMMA matrix. The images have been overlapped with the EDS maps to enlighten the ceramic phase dispersion. It can be observed that the best levels of dispersion were achieved using barium titanate (Fig. 3a) and rutile (Fig. 3c), as expected considering their average size. All three ceramic particles formed clusters, resulting in un-uniform dispersion in the polymeric matrix. Aluminum nitride (Fig. 3b), in particular, resulted in a much coarser surface where partially embedded grains protrude from the matrix.

**Figure 3:**
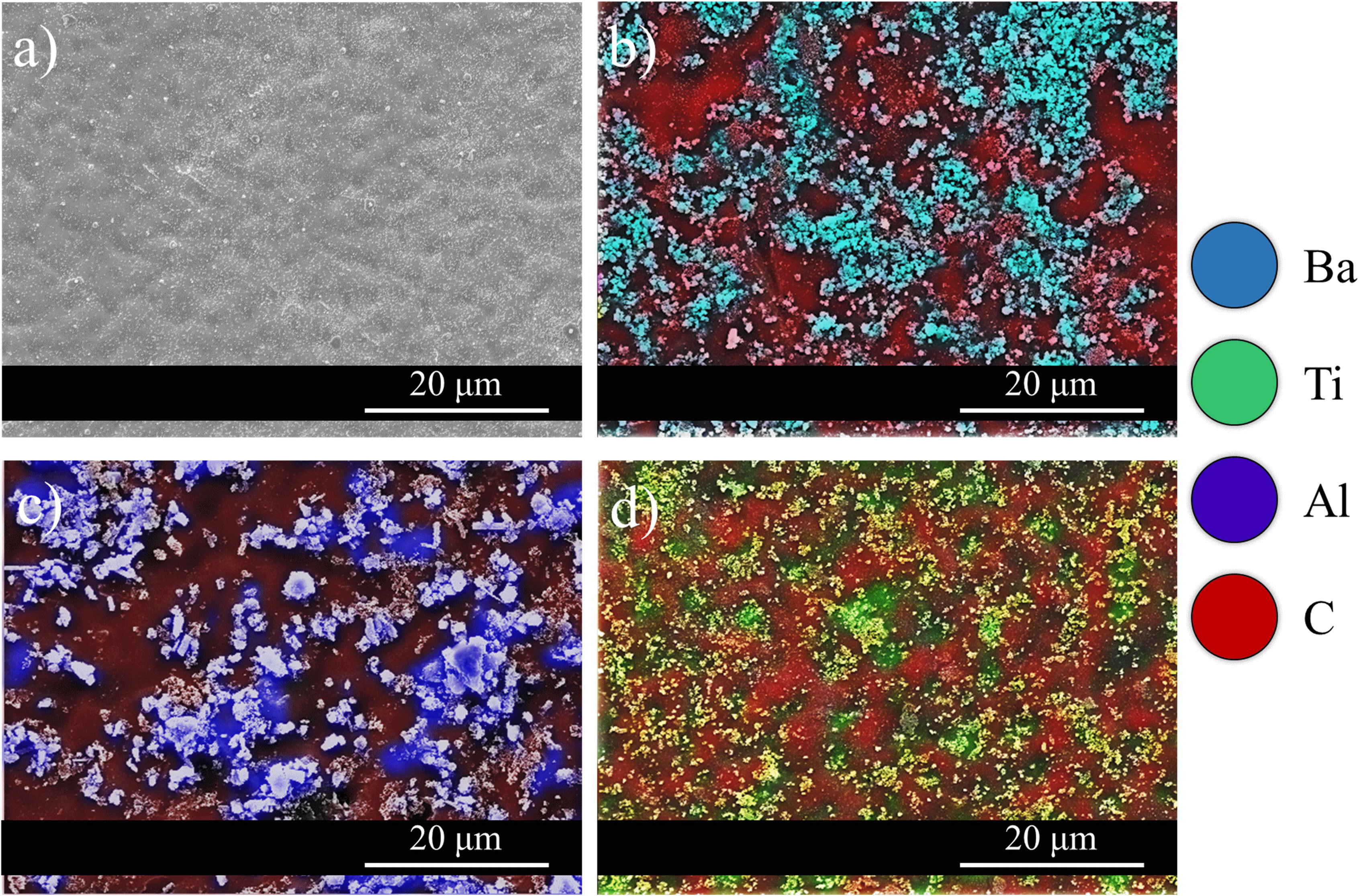
SEM-EDS color maps for different chemical elements, Carbon, Titanium Barium + Titanium and Aluminum. Samples are (a) pristine PMMA, (b) 15% barium titanate, (c) 15% aluminum nitride, (d) 15% rutile

Figure 4 shows the fraction of area containing reinforcing particles with respect to the total investigated area, as detected by SEM-EDS. These results are not directly comparable with the amount of powder introduced for two reasons: the differences in specific volume (6.02 g/cm^3^ for BaTiO_3_, 3.26 g/cm^3^ for AlN and 4.23 g/cm^3^ for TiO_2_, but only 1.18 g/cm^3^ for PMMA) may cause precipitations and the intensity of the X-ray emission measured by SEM-EDS is a function of the chemical element and is usually stronger for heavier elements such as barium when compared to aluminum.

**Figure 4:**
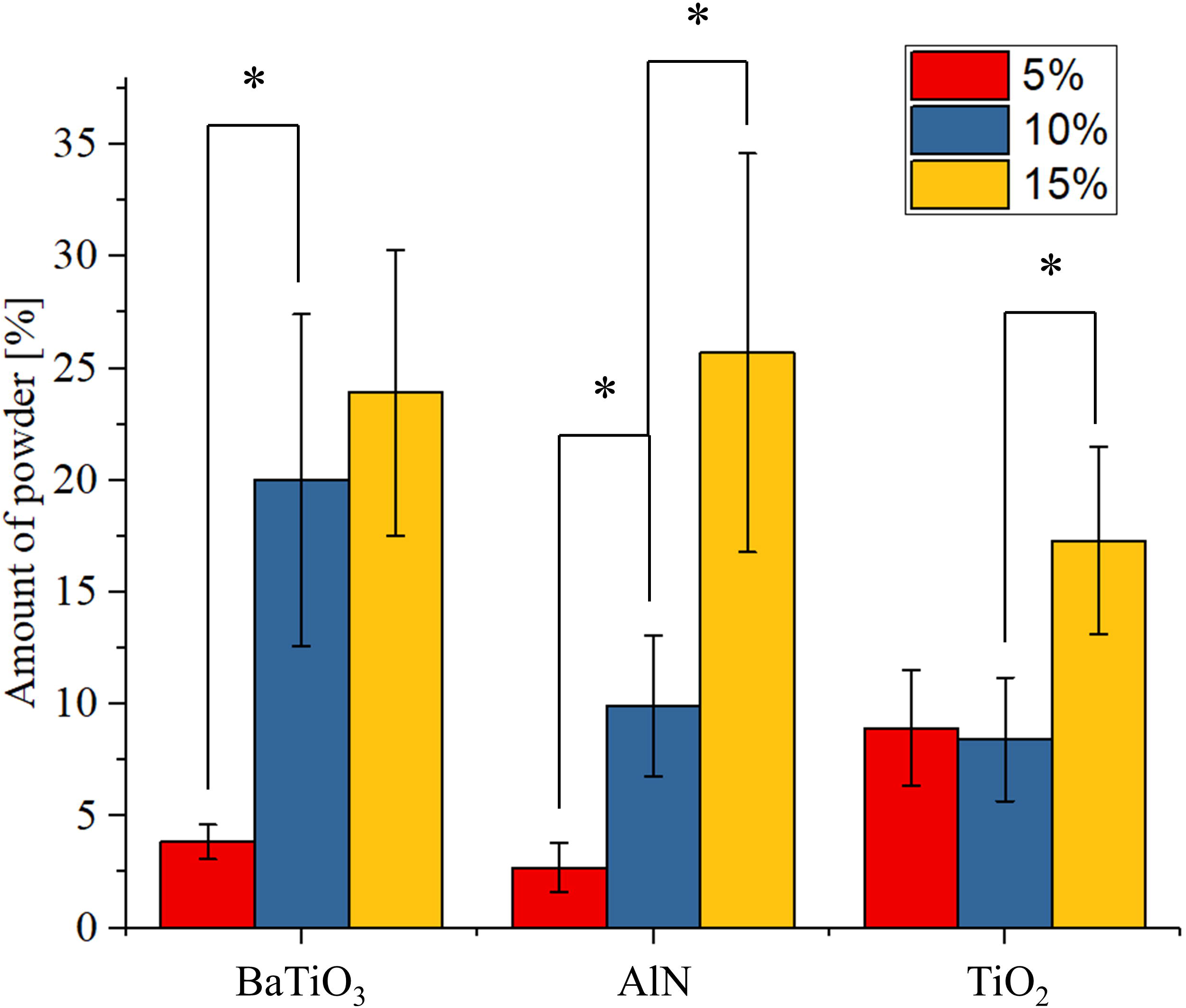
Semi quantitative evaluation of the fraction of area covered by reinforcing particles, for each sample

The large statistical dispersions shown in Fig. 4, as well as the lack of a precise trend between histograms in the same group, indicate that even after a long mixing process the powder are still not homogeneously dispersed. Between the various reinforcing materials, AlN is the only one showing a clear trend when comparing the three concentrations.

Figure 5 shows the quality of the powder dispersion at lower magnifications, as observed by optical/laser microscopy on wider portions of the samples. It can be observed that while powders are unevenly distributed at concentrations of about 5% in weight, at 15% the amount of macroscopic distribution defects has drastically reduced to about 10% of the area or less.

**Figure 5:**
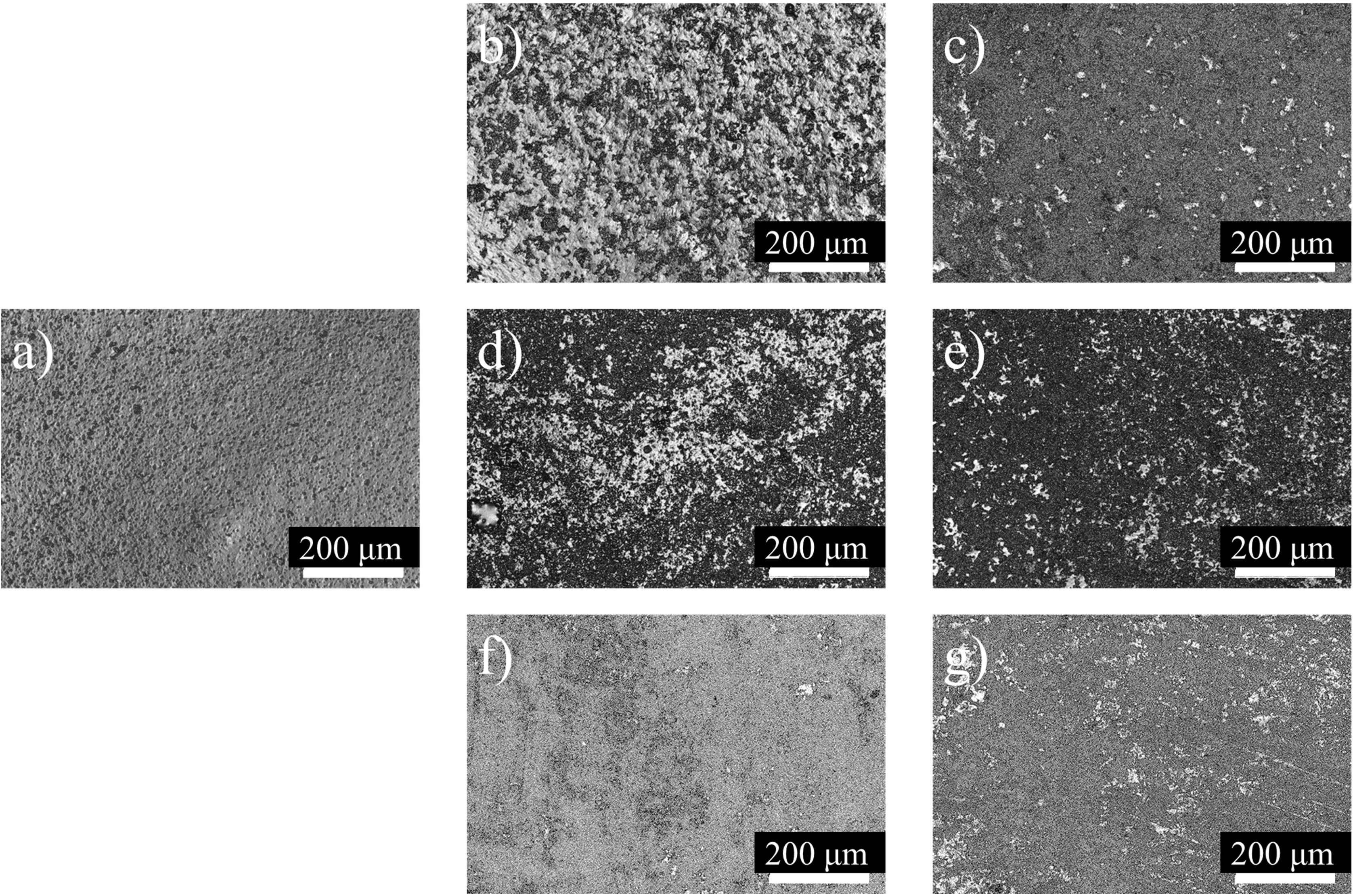
Laser microscope images of the samples surfaces showing the low magnification distribution of the powder particles for samples containing 5% and 15% of reinforcing particles. Samples are (a) pristine PMMA, (b) 5% barium titanate, (c) 15% barium titanate, (d) aluminum nitride (e) 15% aluminum nitride, (f) 5% rutile, (g) 15% rutile

In terms of surface arithmetic roughness (Figure 6), all three ceramic reinforcing particles cause an increase of the arithmetical mean height, proportionally to their concentration. It can be observed that even if the aluminum nitride powder generates the rougher surface at a microscopic level (Fig. 3c), the effect is almost negligible on a larger scale.

**Figure 6:**
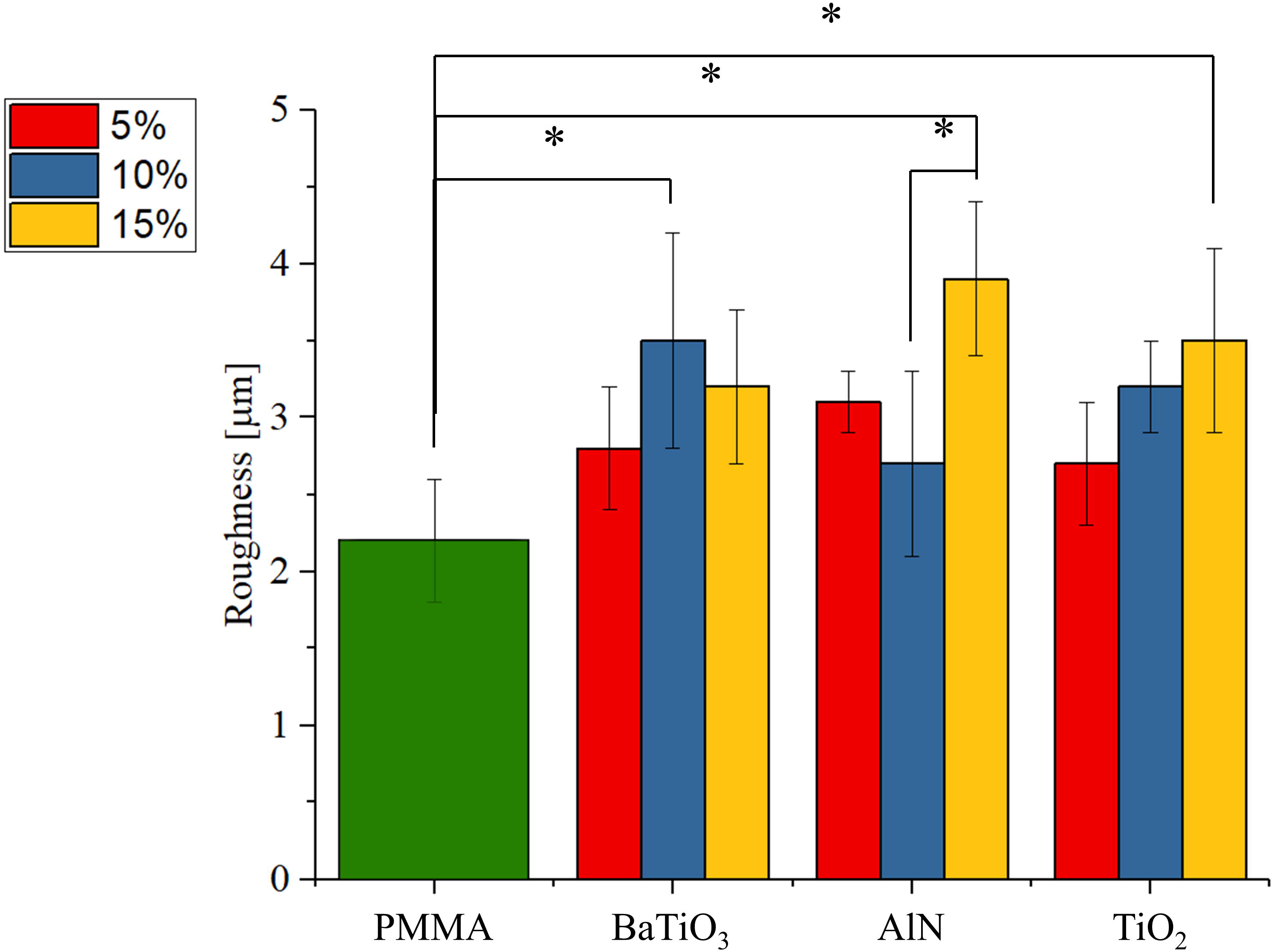
Ra roughness as measured on the different surfaces

Figure 7 shows the Raman spectra acquired on the different samples, as a function of type of reinforcing particles and concentration, as compared to un-reinforced PMMA. The relative Raman bands are listed in Table I, along with literature references for their theoretical assignation [23–27]. It can be observed that the signal coming from the PMMA matrix can be observed on all samples but the ones containing titanium oxide. This is caused by the relatively large Raman cross-section of TiO_2_ when compared to the other reinforcing particles. Moreover, the intensity of the bands associated with the ceramics increase with their concentration, as expected. These results confirm that the particles are well distributed inside the probed volume and even if local differences in intensity were observed from point to point, the average spectra acquired at different regions were similar (r > 0.95). Due to the high transmittance of the laser light (504 nm) in PMMA, Raman results can be considered as representative for the first few hundreds of microns of material.

**Figure 7:**
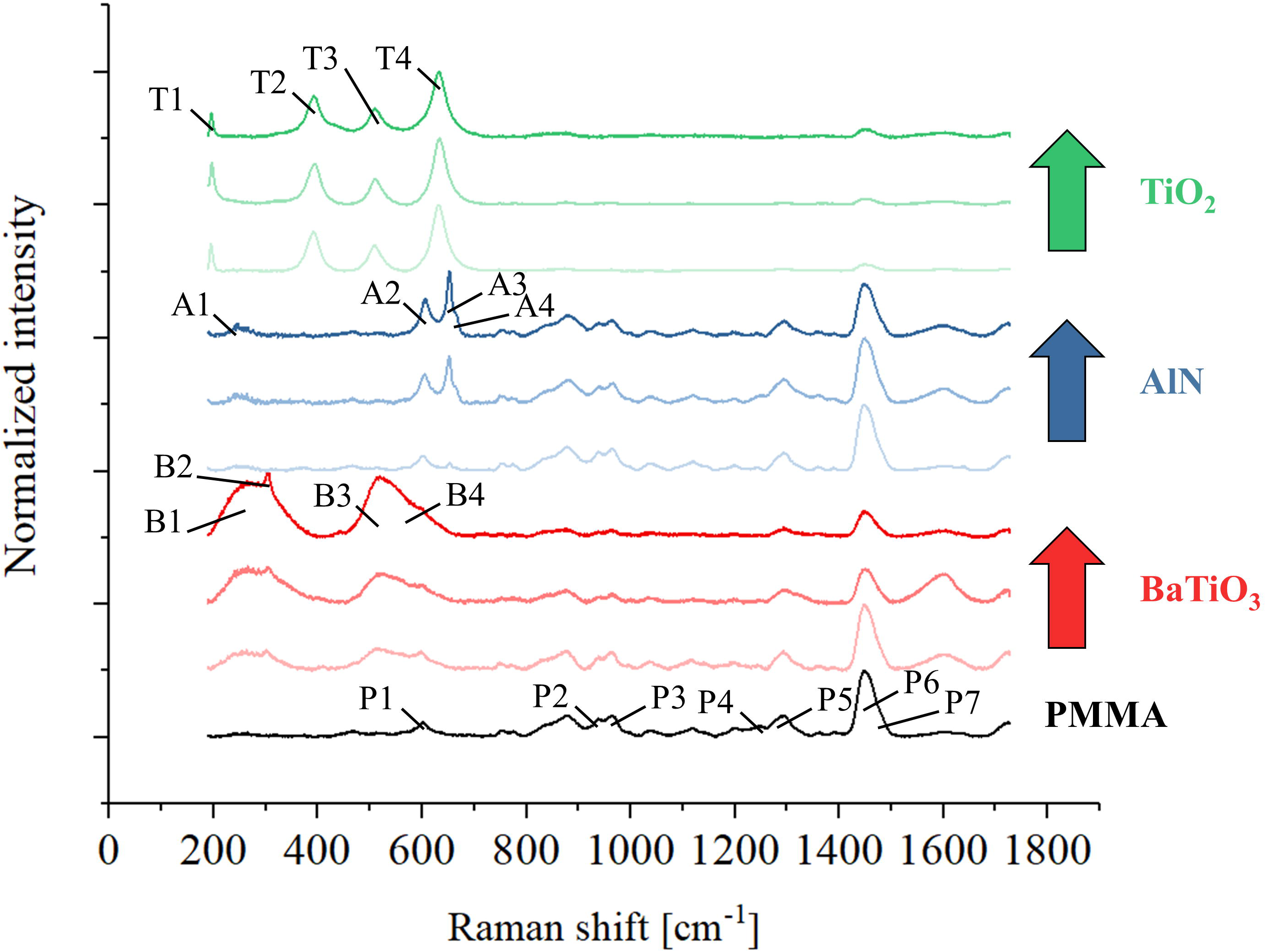
Raman spectroscopy results for all the different samples, as a function of ceramic fraction

**Table I:**
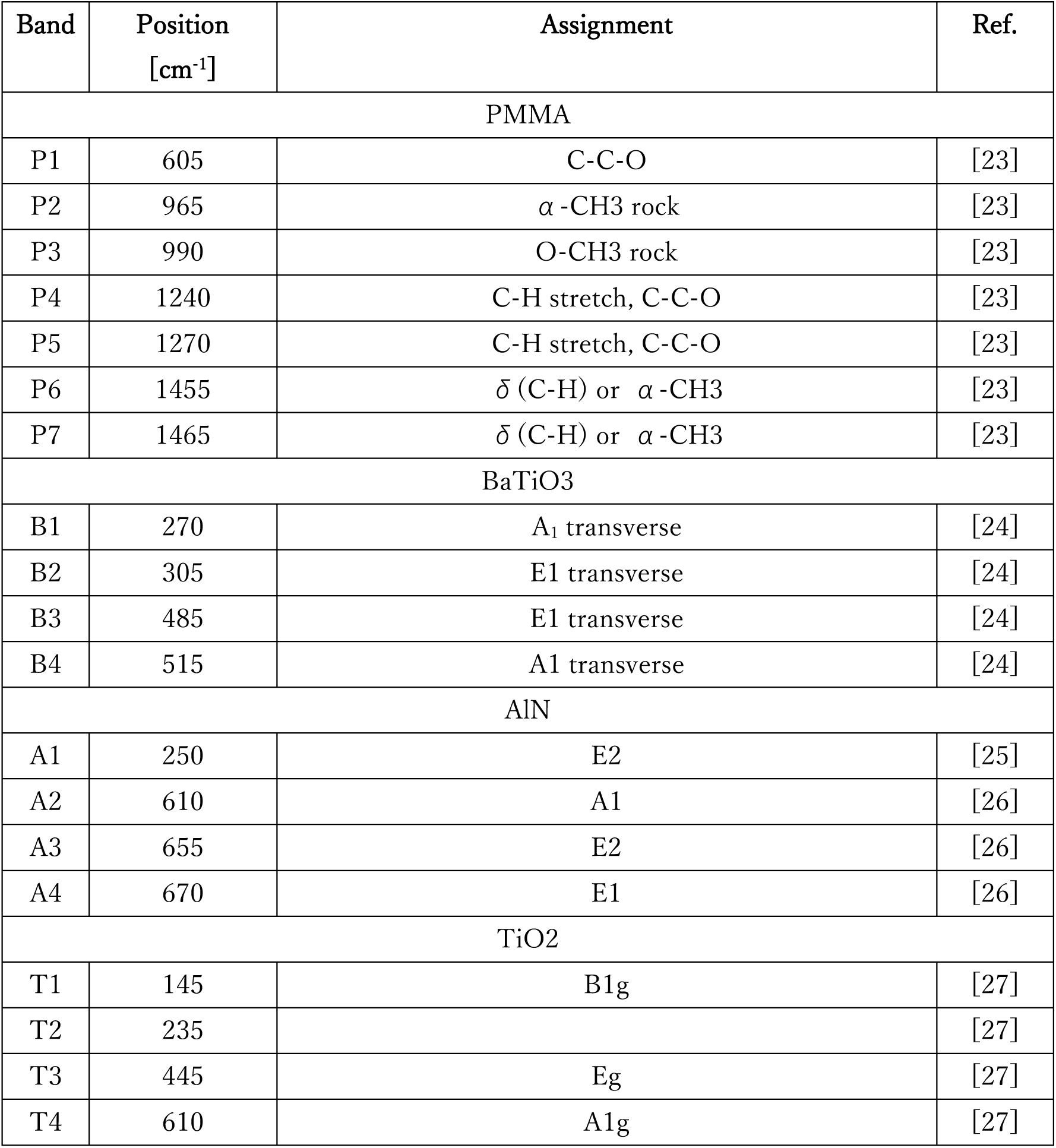
assignation of the Raman bands presented in Figure 7

| Band | Position<br>[cm<sup>-1</sup>] | Assignment | Ref. |
| --- | --- | --- | --- |
| PMMA |  |  |  |
| P1 | 605 | C-C-O | [23] |
| P2 | 965 | $\alpha$ -CH<sub>3</sub> rock | [23] |
| P3 | 990 | O-CH<sub>3</sub> rock | [23] |
| P4 | 1240 | C-H stretch, C-C-O | [23] |
| P5 | 1270 | C-H stretch, C-C-O | [23] |
| P6 | 1455 | $\delta$ (C-H) or $\alpha$ -CH<sub>3</sub> | [23] |
| P7 | 1465 | $\delta$ (C-H) or $\alpha$ -CH<sub>3</sub> | [23] |
| BaTiO<sub>3</sub> |  |  |  |
| B1 | 270 | A<sub>1</sub> transverse | [24] |
| B2 | 305 | E1 transverse | [24] |
| B3 | 485 | E1 transverse | [24] |
| B4 | 515 | A1 transverse | [24] |
| AlN |  |  |  |
| A1 | 250 | E2 | [25] |
| A2 | 610 | A1 | [26] |
| A3 | 655 | E2 | [26] |
| A4 | 670 | E1 | [26] |
| TiO<sub>2</sub> |  |  |  |
| T1 | 145 | B1g | [27] |
| T2 | 235 |  | [27] |
| T3 | 445 | Eg | [27] |
| T4 | 610 | A1g | [27] |

Figure 8 shows the FTIR spectra acquired on the different samples, as a function of type of reinforcing particles and concentration, as compared to un-reinforced PMMA. The relative FTIR bands are listed in Table II, along with literature references [28–31]. When compared to Raman, the FTIR probe is shallow, representing only the regions closer to the external surface. Moreover, the probe is larger than the diameter of the Raman and so less affected by micrometric range powder distributions. Unlike Raman, the FTIR spectra of PMMA is more intense and sharper than those of the reinforcing particles, but the dependence of concentration can still be clearly observed for all three materials, for example by comparing the region between 450 and 1000 cm^-1^ with the intense band at about 1700 cm^-1^.

**Figure 8:**
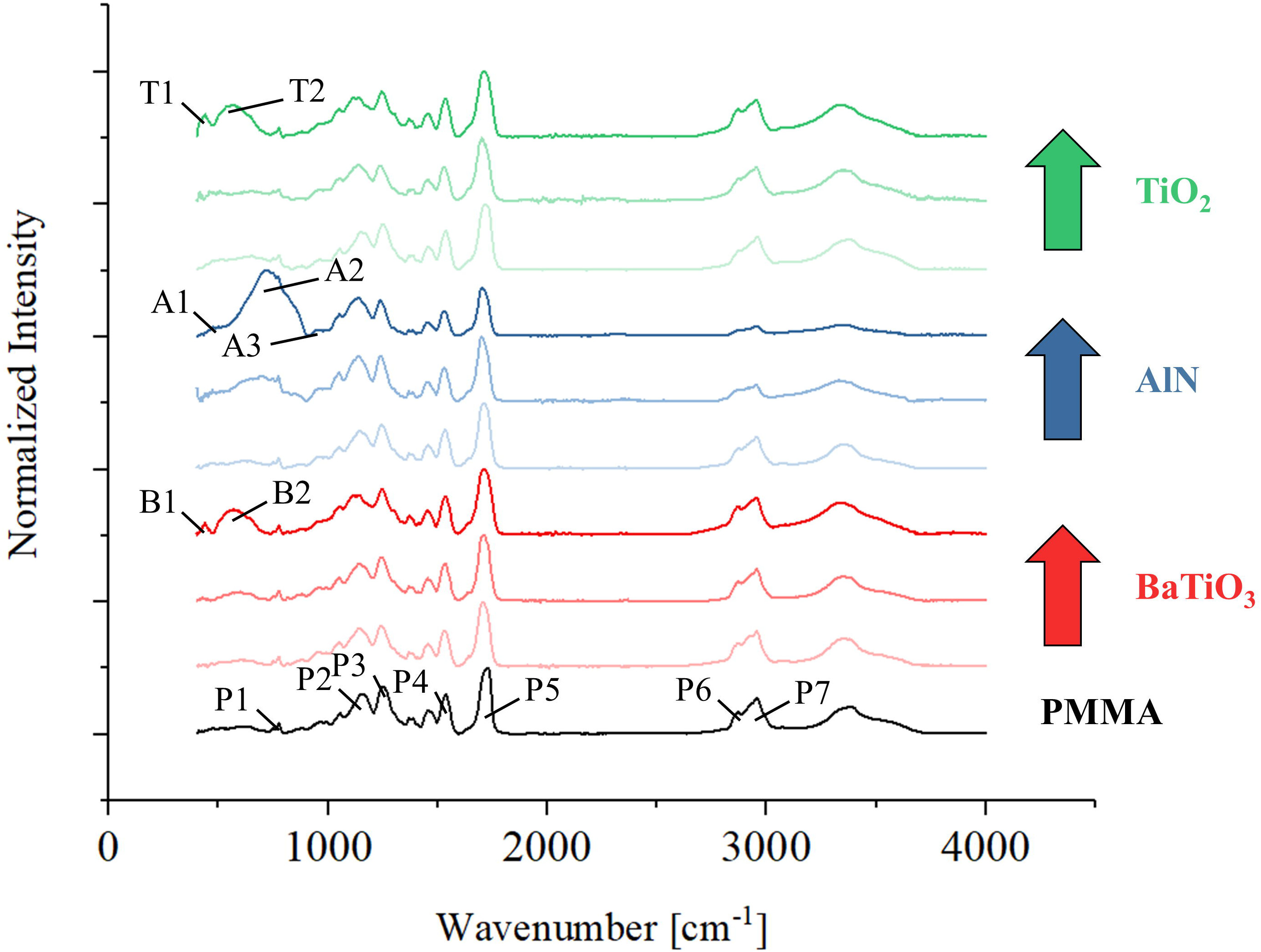
FTIR spectroscopy results for all the different samples, as a function of ceramic fraction

**Table II:**
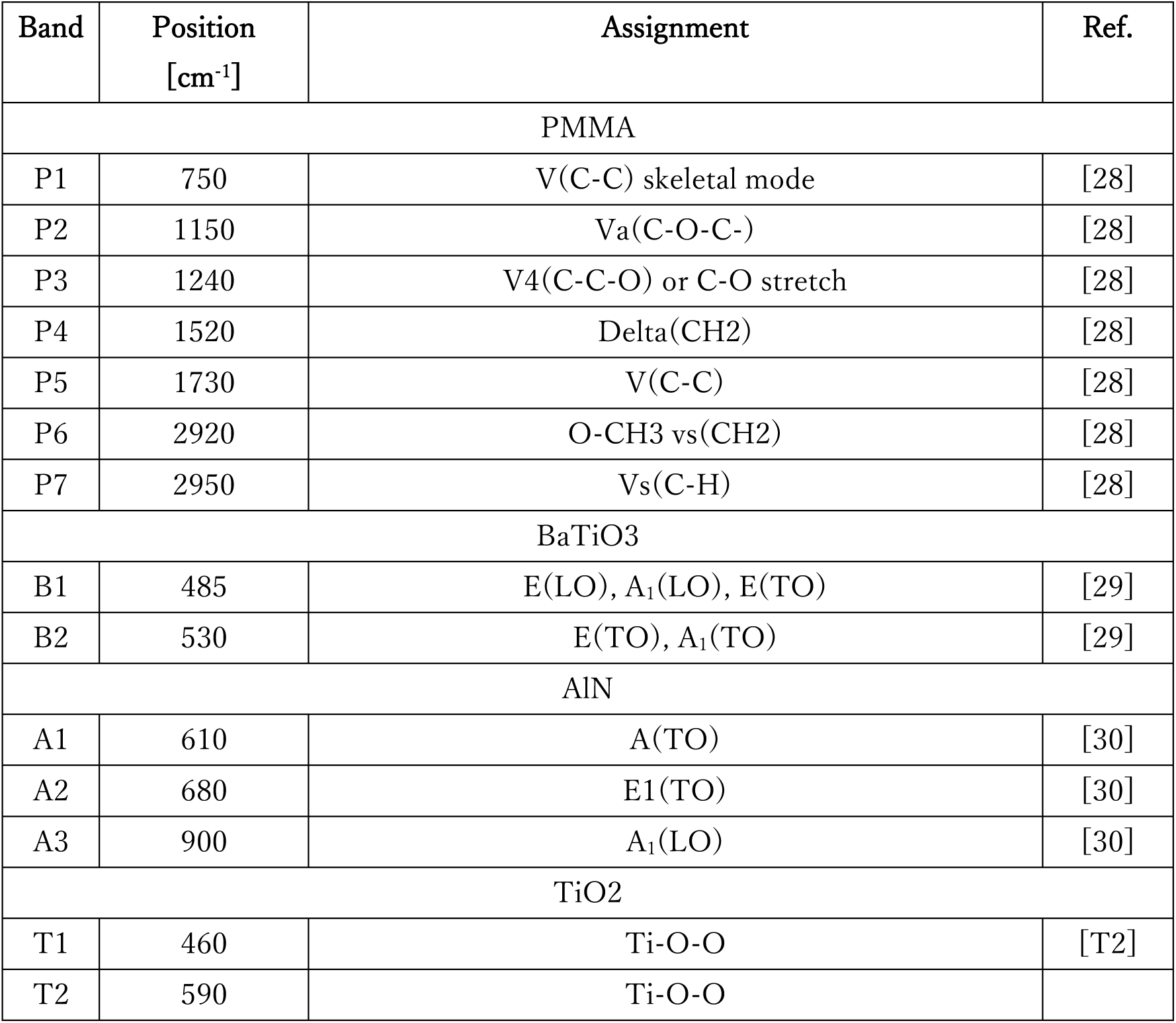
assignation of the FTIR bands presented in Figure 8

Figure 9 shows the results of the colony counting unit test performed on the samples with 5 and 15% of ceramic powders. It can be observed that both BaTiO_3_ and AlN show a reduction on the number of bacteria both after 24 and 48 hours of treatment. TiO_2_, on the other hand, shows a comparable amount of bacteria respect to the pure PMMA reference. The two antibacterial particles seem to have a different effect over time: while BaTiO_3_ is more effective at 24 hours of treatment, AlN results are lower at 48 hours.

**Figure 9:**
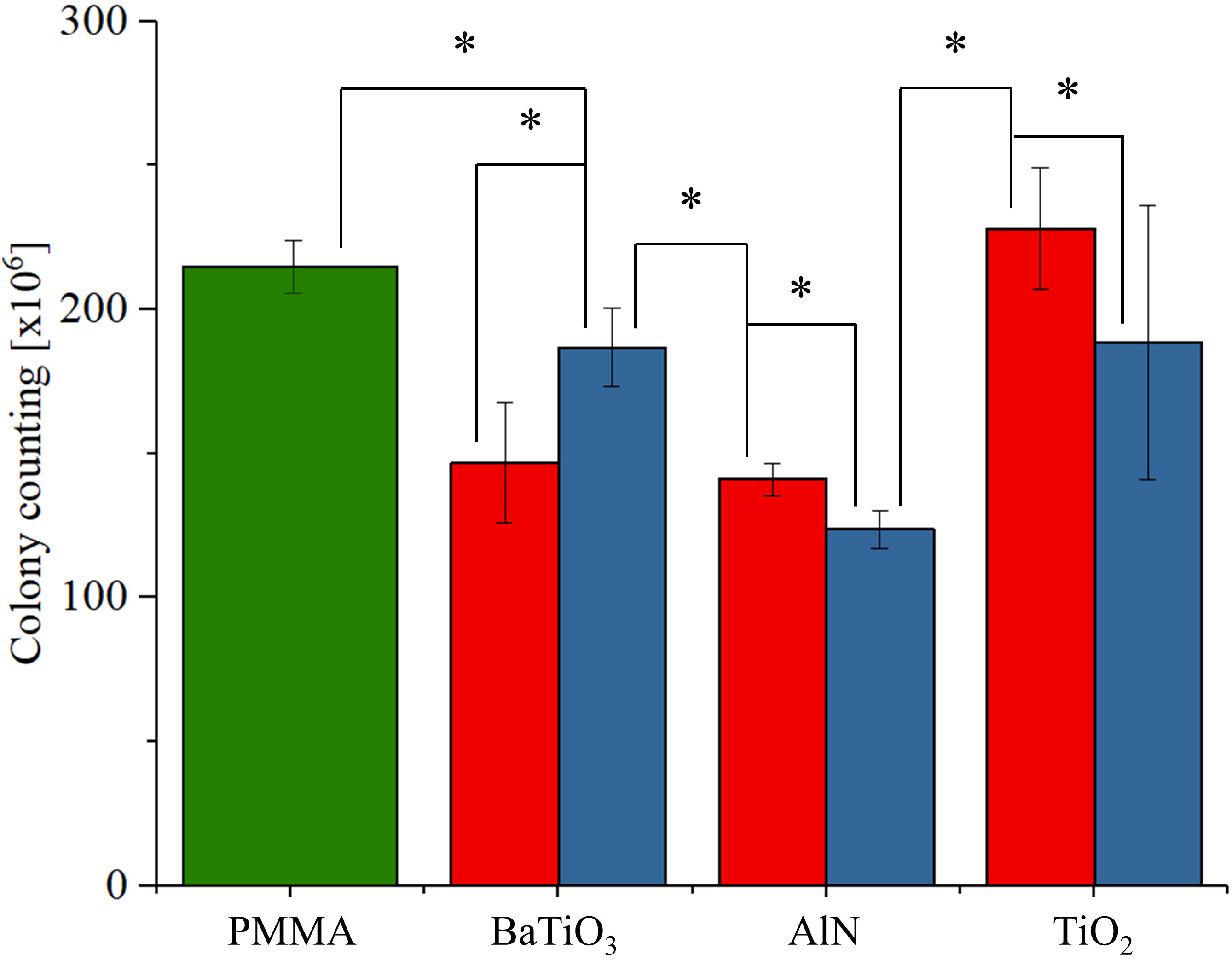
Colony counting units as measured after 24 and 48 hours of *in vivo* testing with *E. coli*

Figure 10 shows the results of the Raman imaging performed on the PMMA resin containing 15% of AlN and 3D printed into a construction block using stereo-lithography. Purple areas are mainly constituted by PMMA matrix while the yellow particles are aluminum nitride powders. The relative spectral regions are marked in the inlet on the right . It can be observed that the ceramic reinforcing particles are well distributed in the matrix, with no noticeable defect produced by the printing process or the polymer curing.

**Figure 10:**
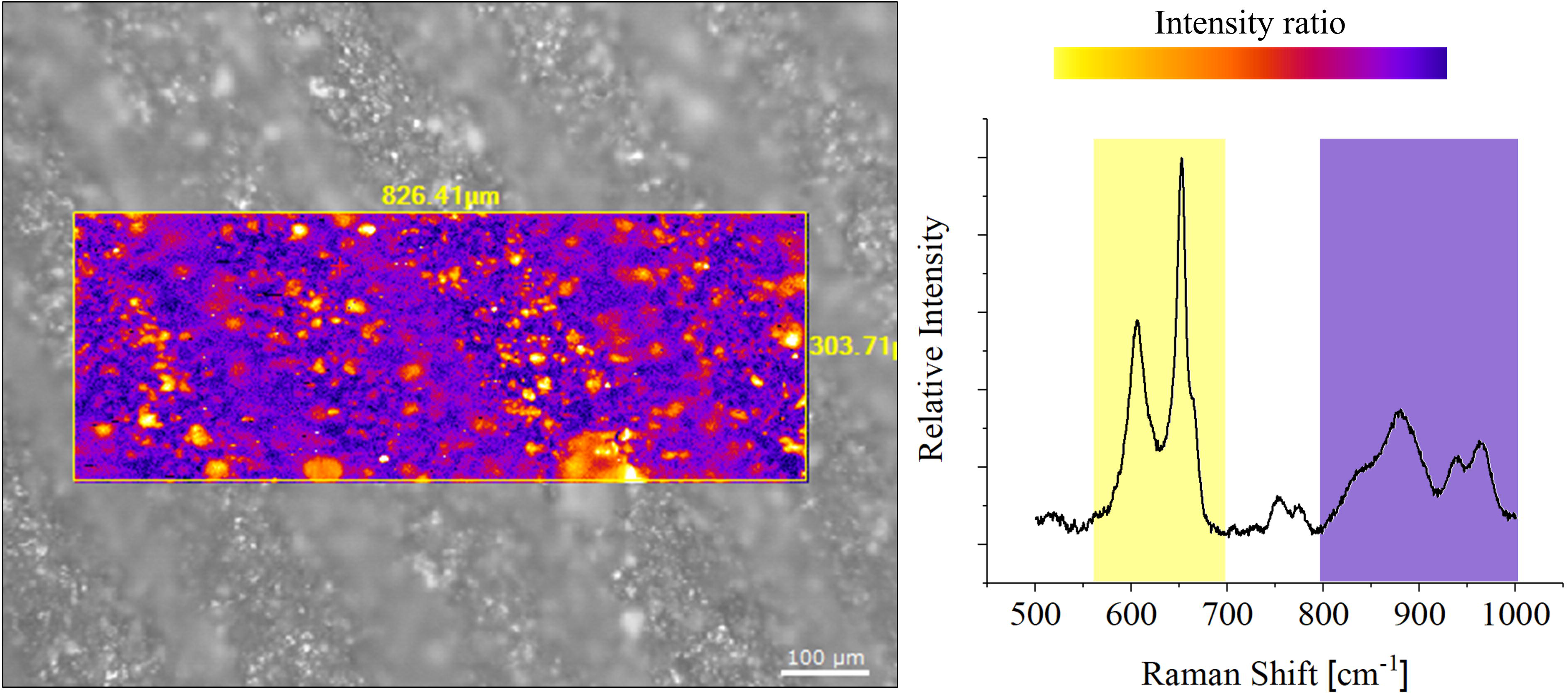
Raman imaging showing the distribution of aluminum nitride powders inside the PMMA matrix, for a sample containing 15% of reinforcing ceramic particles

Figure 11 shows the results of the mechanical testing performed on the 3D printed specimen containing different concentrations of AlN powder. Mechanical properties decrease with increasing the amount of ceramic, both in the as-printed and in the post-cured conditions. In the post-cured conditions, the mechanical strength is about 120% higher than after 3D printing, but the addition of about 15% of AlN reduces the ultimate strength to about 80% the value obtained on pure PMMA. The addition of 15% of AlN reduced the ultimate strain from about 4.2% to 3.3% for not UV cured samples and from 3.2% to 1.4% for cured samples, meaning that the samples become brittle.

**Figure 11:**
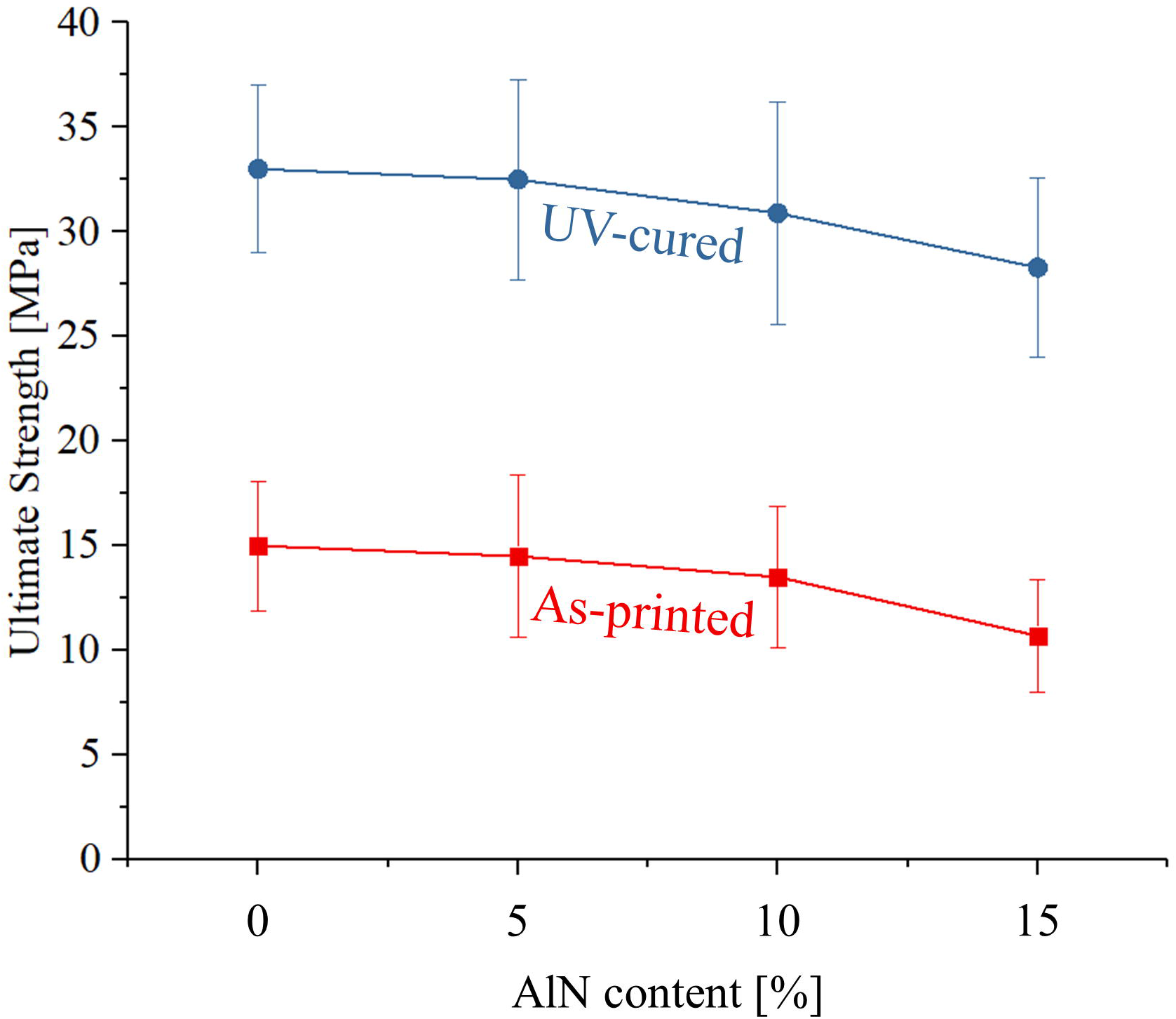
mechanical testing results (ultimate strength) as obtained on samples containing variable fractions of aluminum nitride powders, before and after UV curing post-treatment.

Figure 12 shows the results of the WST testing performed on the 3D printed parts containing 5% and 15% of aluminum nitride, compared with the results from the pristine PMMA part. It can be observed that the amount of bacteria on the surface of the samples drastically decreases with increasing the fraction of reinforcing particles and exposure time. Samples containing 15% of aluminum nitride treated with bacteria for 48 hours have a statistically significant lower optical density when compared to the same samples at 24 hours, while the opposite behavior is shown on the pristine material.

**Figure 12:**
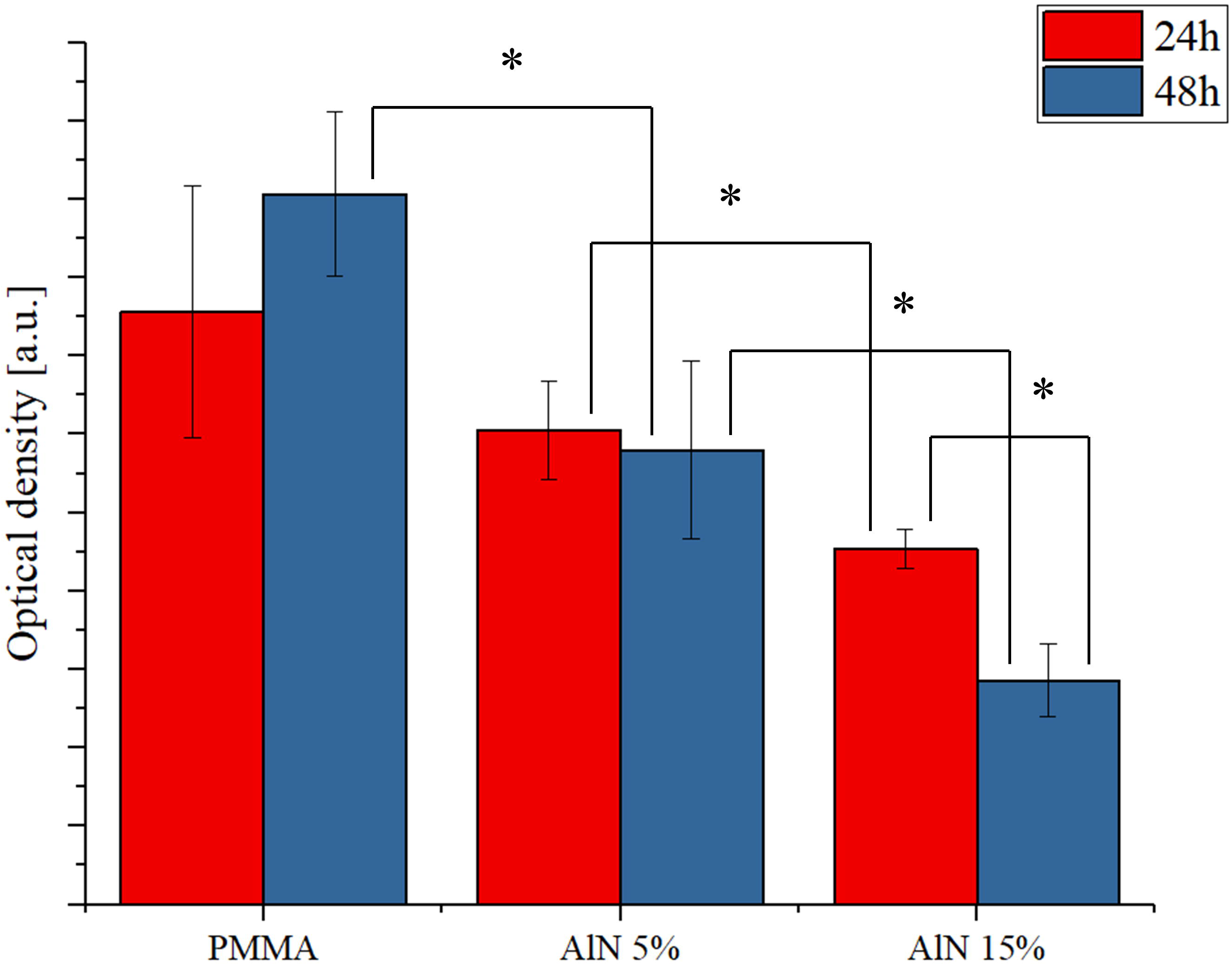
Results of the WST testing after *in vitro* testing with *E. coli* on the 3D printed samples containing 5 and 15% of aluminum nitride, compared to the pristine PMMA material

Figure 13 shows the results of the luciferin testing performed on the complex 3D blocks printed using stereo-lithography. It can be observed that the sample containing aluminum nitride has a decrease in both luminescent areas and luminescence intensity, meaning that these surfaces are resistant to bacteria colonization or actively anti-bacterial. The composite containing 5% of reinforcing particles shows the presence of a few residual spots with bacteria around the edges, while the the sample containing 15% of aluminum nitride there is no visible bacteria colonization.

**Figure 13:**
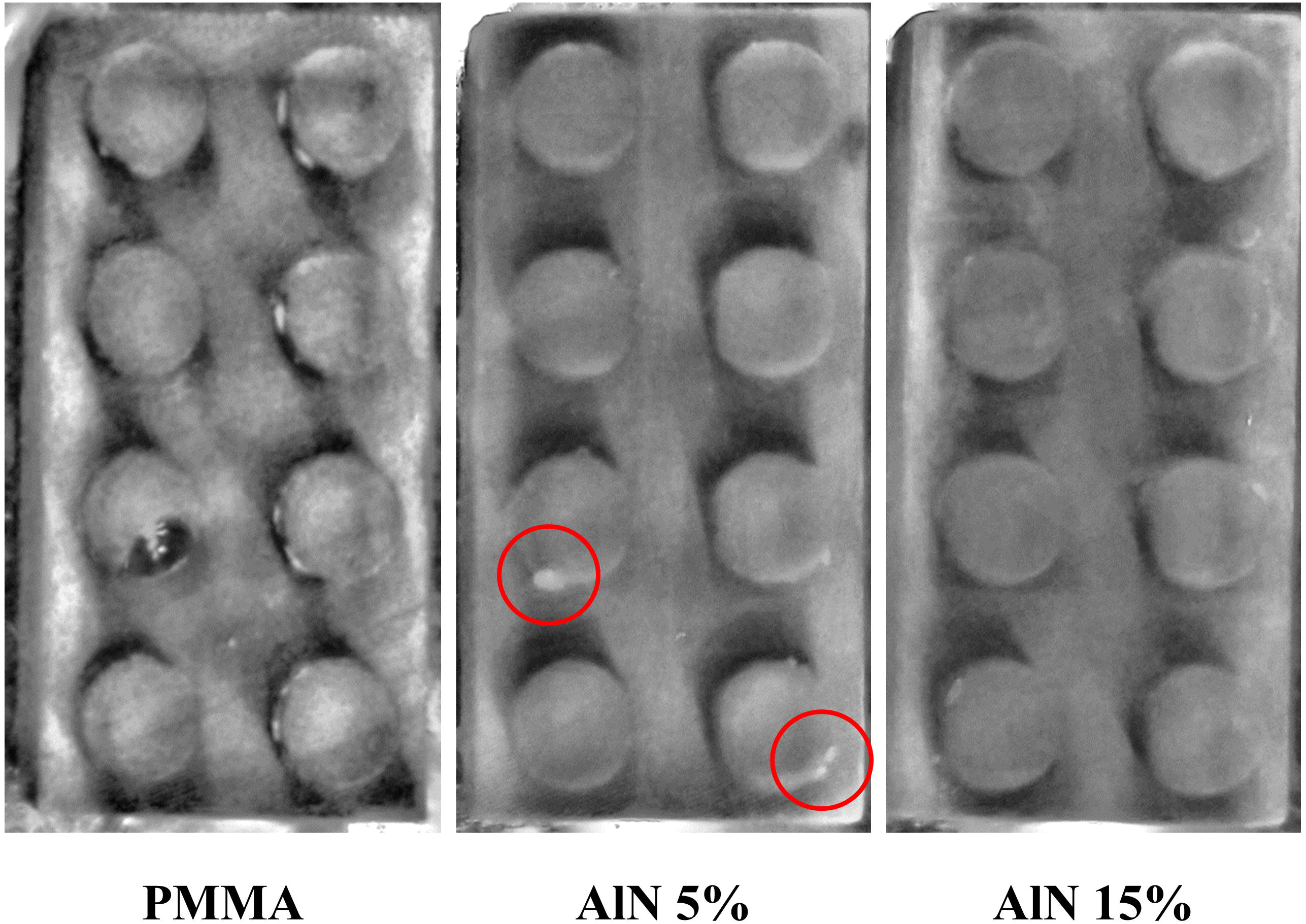
UV lamp detection of residual bacterial matter on the surface of complex 3D printed composite parts: (a) pristine PMMA, (b) 5% aluminum nitride and (c) 15% aluminum nitride

## 4. Discussion

Effective, polymer-based antibacterial composite materials reinforced with barium titanate and aluminum nitride were already successfully developed in previous studies [19–21].

While barium reinforced Polyvinyl-siloxane scaffolds were obtained by a conventional vacuum casting technique [19], previous attempts to use AlN in 3D printing were limited to a final spray-coating layer of resin mixed with reinforcing particles. To print resin composites using stereo-lithography can prove to be a difficult challenge as even low amounts of reinforcement phases can drastically modify the optical parameters and in particular the light transmittance.

Rutile-based composites were discarded at a preliminary stage due to the lack of antibacterial efficacy and stereolithographic 3D printing of barium titanate composites was unsuccessful. Powders agglomerated during processing resulting in irregular surfaces and physical discontinuities in the final pieces. Moreover, during processing of large surfaces, a thick layer of barium titanate-enriched polymer formed on the printer base, reducing the efficiency of the laser beam.

It has been postulated that the smaller diameter of the reinforcing powder is responsible for the difficulties during processing, in particular for high fraction of titanate.

When AlN powders were used, the printing resulted in a surface finishing comparable to that of pristine PMMA resin. The slight increase in surface roughness caused by the presence of particles resulted to be about one order of magnitude smaller than the layer resolution of the 3D printer. This can be clearly observed in Fig. 10: the average diameter of the yellow particles in the Raman imaging map is much smaller than the distance between two consecutive melted lines (black arrows), while the depth of field of the optical image is unaffected by the local presence of reinforcements.

Surprisingly, the ceramic reinforcing powder was homogeneously dispersed in the resin during the overall printing, without applying any additional dispersion system such as an ultrasonic sonicator. This was thanks to the similarly negative charge associated with both the AlN powder [32] and the PMMA polymer [33]. Strongly charged particles are conventionally considered easier to disperse, and both PMMA and AlN are polar [34] with the AlN powders being relatively far from the isoelectric point [32] at the pH conditions of the resin [35].

Taking into consideration the antibacterial properties, the results of this research are in line with previously published data [20]: PMMA AlN composites exhibit a clear antibacterial effect against *E. coli*.

## 5. Conclusions

In this research, PMMA stereo-lithographic resins were combined with three different ceramic reinforcing particles and then in vitro tested with *E. coli* bacteria.

Both barium titanate and aluminum nitride composites resulted to be antibacterial, with latter showing the overall best performances.

When 3D printed as a composite, aluminum nitride reinforced PMMA showed a good phase distribution and geometrical precision, comparable to the pristine material.

3D printed composites obtained with aluminum nitride resulted to be resistant to colonization from *E coli*.

## Data availability statement

Data available on request from the authors.

