## Supplementary material for "Antibacterial 3D-printed PMMA/ceramic composites": Table I

Table I: assignation of the Raman bands presented in Figure 7

| **Band** | **Position**  **[cm^-1^]** | **Assignment** | **Ref.** |
| --- | --- | --- | --- |
| PMMA | | | |
| P1 | 605 | C-C-O | [23] |
| P2 | 965 | α-CH3 rock | [23] |
| P3 | 990 | O-CH3 rock | [23] |
| P4 | 1240 | C-H stretch, C-C-O | [23] |
| P5 | 1270 | C-H stretch, C-C-O | [23] |
| P6 | 1455 | δ(C-H) or α-CH3 | [23] |
| P7 | 1465 | δ(C-H) or α-CH3 | [23] |
| BaTiO3 | | | |
| B1 | 270 | A_1_ transverse | [24] |
| B2 | 305 | E1 transverse | [24] |
| B3 | 485 | E1 transverse | [24] |
| B4 | 515 | A1 transverse | [24] |
| AlN | | | |
| A1 | 250 | E2 | [25] |
| A2 | 610 | A1 | [26] |
| A3 | 655 | E2 | [26] |
| A4 | 670 | E1 | [26] |
| TiO2 | | | |
| T1 | 145 | B1g | [27] |
| T2 | 235 |  | [27] |
| T3 | 445 | Eg | [27] |
| T4 | 610 | A1g | [27] |
