## Supplementary material for "Antibacterial 3D-printed PMMA/ceramic composites": Table II

Table II: assignation of the FTIR bands presented in Figure 8

| **Band** | **Position**  **[cm^-1^]** | **Assignment** | **Ref.** |
| --- | --- | --- | --- |
| PMMA | | | |
| P1 | 750 | V(C-C) skeletal mode | [28] |
| P2 | 1150 | Va(C-O-C-) | [28] |
| P3 | 1240 | V4(C-C-O) or C-O stretch | [28] |
| P4 | 1520 | Delta(CH2) | [28] |
| P5 | 1730 | V(C-C) | [28] |
| P6 | 2920 | O-CH3 vs(CH2) | [28] |
| P7 | 2950 | Vs(C-H) | [28] |
| BaTiO3 | | | |
| B1 | 485 | E(LO), A_1_(LO), E(TO) | [29] |
| B2 | 530 | E(TO), A_1_(TO) | [29] |
| AlN | | | |
| A1 | 610 | A(TO) | [30] |
| A2 | 680 | E1(TO) | [30] |
| A3 | 900 | A_1_(LO) | [30] |
| TiO2 | | | |
| T1 | 460 | Ti-O-O | [T2] |
| T2 | 590 | Ti-O-O |  |
